# Alterations of metagenomics and metaproteomics associate kidney disease in a combination of opisthorchiasis and nonalcoholic fatty liver disease

**DOI:** 10.1101/2023.09.20.558740

**Authors:** Keerapach Tunbenjasiri, Thasanapong Pongking, Chutima Sitthirach, Suppakrit Kongsintaweesuk, Sitiruk Roytrakul, Sawanya Charoenlappanit, Sirinapha Klungsaeng, Sirirat Anutrakulchai, Chalongchai Chalermwat, Somchai Pinlaor, Porntip Pinlaor

## Abstract

**Background:** Non–alcoholic fatty liver disease (NAFLD) is prevalent worldwide and is associated with chronic kidney disease (CKD). *Opisthorchis viverrini* (Ov) infection and consumption of high-fat and high-fructose (HFF) diets exacerbate NAFLD leading to nonalcoholic steatohepatitis. Here, we aimed to investigate the effects of a combination of HFF diets and *O.viverrini* infection on kidney pathology via changes in the gut microbiome and host proteome in hamsters.

**Methodology/Principal findings:** Twenty animals were divided into four groups; Normal diet feeding and non-infected Ov (Normal); HFF diets feeding (HFF); Ov infection (Ov); and feeding with a combination of HFF diets and infection with Ov (HFFOv). Fecal samples were extracted and used for Illumina Miseq sequencing platform based on the V3–V4 region of the 16S rRNA gene, along with LC/MS-MS analysis. Histopathological studies and biochemical assays were also conducted. The results indicated that the HFFOv group exhibited the most severe kidney injury, as elevated KIM-1 expression and accumulation of fibrosis in kidney tissue. In comparison with the HFF group, the combined group showed higher diversity and composition. An increased number of *Ruminococaceae*, *Lachospiraceae*, *Desulfovibrionaceae* and *Akkermansiaceae*, and a lower number of *Eggerthellaceae* were observed. A total of 243 significant host proteome were identified in all groups. Analysis using STITCH predicted that host proteome associated leaky gut such as soluble CD14 and p-cresol may play a role in the development of kidney disease. Among host proteome, TGF-beta, involving in fibrogenesis, was significantly expressed in HFFOv.

**Conclusions/Significance:** The combination of HFF diets and *O.viverrini* infection may promote kidney injury through the alterations in the gut microbiome and host proteome. This knowledge may be an effective strategy to prevent the progression of CKD beyond the early stages.

**Author summary:** A diets high in fat and fructose causes nonalcoholic fatty liver disease (NAFD), which is increasing worldwide. Liver fluke (*Opisthorchis viverrini*, Ov) infection is endemic in the Mekong subregion including in the northeastern Thailand. The prevalence of opisthorchiasis caused by the infection with *O. viverrini* is associated with fatty liver and bile duct cancer. We have previously demonstrated that infection with *O. viverrini* exacerbates NAFD progression to non-alcoholic steatohepatitis (NASH) in animal models. NASH exists kidney disease severity higher than ingestion of high-fat and high-fructose (HFF) diets or infection with *O. viverrini*. Here, we further investigate whether metagenomics is more likely to change in NASH than in NAFD or opisthorchiasis conditions. The combined group had higher diversity and composition. Elevated levels of *Ruminococaceae*, *Lachospiraceae*, *Desulfovibrionaceae* and *Akkermansiaceae* and decreased levels of *Eggerthellaceae* were observed, suggesting that HFF+Ov may cause gut dysbiosis in NASH. Differentially expressed proteins were also associated with these gut microbiomes in NASH condition. In addition, we found that the association of metagenomics and metaproteomics in NASH was related to kidney disease. Analysis using STITCH predicted that host proteome may be involved in leaky gut such as soluble CD14 and p-cresol to promote kidney disease. A significantly expressed TGF-beta involving fibrogenesis was found to be associated with kidney fibrosis. Therefore, alterations of metagenomics and metaproteomics is associated with kidney disease in a combination of opisthorchiasis and nonalcoholic fatty liver disease.

## Introduction

Fatty liver disease is divided into 2 major types depending on alcohol consumption [1]. One fatty liver disease, non–alcoholic fatty liver diseases (NAFLD), refers to fat accumulation in the liver that is not due to alcohol consumption. The worldwide prevalence of NAFLD is reported to be approximately 20% to 50% [2]. The high fat, high sugar dietary pattern was associated with incidence of chronic kidney disease (CKD) [3]. In addition, *Opisthorchis viverrini* (Ov) infection is common and affects at least six million people, especially in the northeast Thailand. Ultrasonography screening in Khon Kaen found *O. viverrini* infection in 10% of patients with fatty liver disease[4, 5]. Chronic *O. viverrini* infection was associated with kidney pathology in the patients [6]. Long-term intake of high-fat and high-fructose (HFF) diets [7, 8] or chronic *O. viverrini* infection [9] can cause renal pathology in animal models. The severity of kidney pathology is more likely severe in the condition of two factors than each agent [10]. Thus, understand of the underlying mechanism of this combination risk factor of kidney disease would be benefit for prevention CKD progression, particularly in endemic area of infectious agents.

Chronic kidney disease (CKD) is defined as abnormalities in both the structure and/or function of kidney. The increasing incidence and prevalence of CKD are major health concerns, impacting approximately 10% of the global population [11]. In Thailand, there is a high prevalence of 17.5% was reported in 2009, with the highest at 22.2% in the northeastern region. CKD is believed to be a multifactorial disease. Elderly individuals, genders, racial minorities, females, and those with diabetes mellitus and hypertension are all associated with a higher prevalence of CKD [12]. Moreover, either infection with *O. viverrine* [13, 14] or *Strongyloides stercoralis*, consumption of either monosodium glutamate (MSG) or high-fat and high-fructose (HFF) diets change the composition of the gut microbiome called gut dysbiosis, thus contributing to the development of various diseases including CKD [15, 16]. A previous study revealed that a combination of *O. viverrini* infection and HFF diets consumption causes liver and kidney pathology [17]. However, a combination of HFF diets and *O. viverrini* infection causes CKD is still unclear, particularly via the alterations in gut microbiota and proteomics.

Gut microbial alterations, which are called gut dysbiosis, are a key role in many inflammatory-related diseases. It influences the permeability of the intestinal mucosal barrier and releases inflammatory factors and endotoxins in the bloodstream [18–20]. Many previous studies reported gut microbiome change associated CKD both of human and animal [16, 21]. In addition, Liquid chromatography tandem mass spectrometry (LC-MS/MS) technologies have the capacity to provide deeper information for both identification and quantification of proteins in fecal samples[22, 23]. Proteomics-based approaches are used for improving understanding of the physio pathological mechanisms of CKD and identification of new CKD-related biomarkers[24]. Using these two combinations of metagenomic and proteomics technologies would explore the pathophysiology of renal dysfunction and insights into treatment and prevention of CKD [25, 26].

In this study, we hypothesized that a combination of *O.viverrini* infection and consumption of HFF diets promotes the kidney injury through the gut microbiome changes. In addition, we investigated metaproteomics for identification of biomarkers that associate kidney injury. The result from this study may be an effective strategy to prevent the progression of CKD.

## Materials and Methods

### Ethic statement

The samples in this study were used from a previous study (IACUC–KKU–81/62) [27]. All of the experimental protocols were reviewed and approved by the Animal Ethic Committee of Khon Kaen University (IACUC–KKU–88/65). Hamsters were obtained from the Animal Unit, Faculty of Medicine, Khon Kaen University.

### Experimental design

Twenty male Syrian golden hamsters (*Mesocricetus auratus*: 4–6 weeks old and 80–100 g) were used for the experiment. Hamsters were divided into 4 groups (n = 5 each) and animals were sacrificed at the end of 3 months. The name list for each group: normal control (Normal: non-fed high fat and high fructose (HFF) diets, and non-infected *Opisthorchis viverrini* (Ov)); hamsters fed only HFF diets; hamsters fed only *O. viverrini* infection (Ov) and hamsters fed HFF diets and infected with *O. viverrini* (HFFOv). Serum and urine sample were stored at -20 °C until use to determine biochemical kidney function. A portion of each kidney sample was preserved 10% formalin for histopathological and immunohistochemical analysis. Fecal sample was snap frozen in liquid nitrogen for 16S-rRNA next-generation sequencing (16S-rRNA NGS) and Liquid chromatography with tandem mass spectrometry (LC-MS/MS) for gut microbiome and proteomic) analysis respectively as shown in (**Fig 1).**

**Fig 1.**
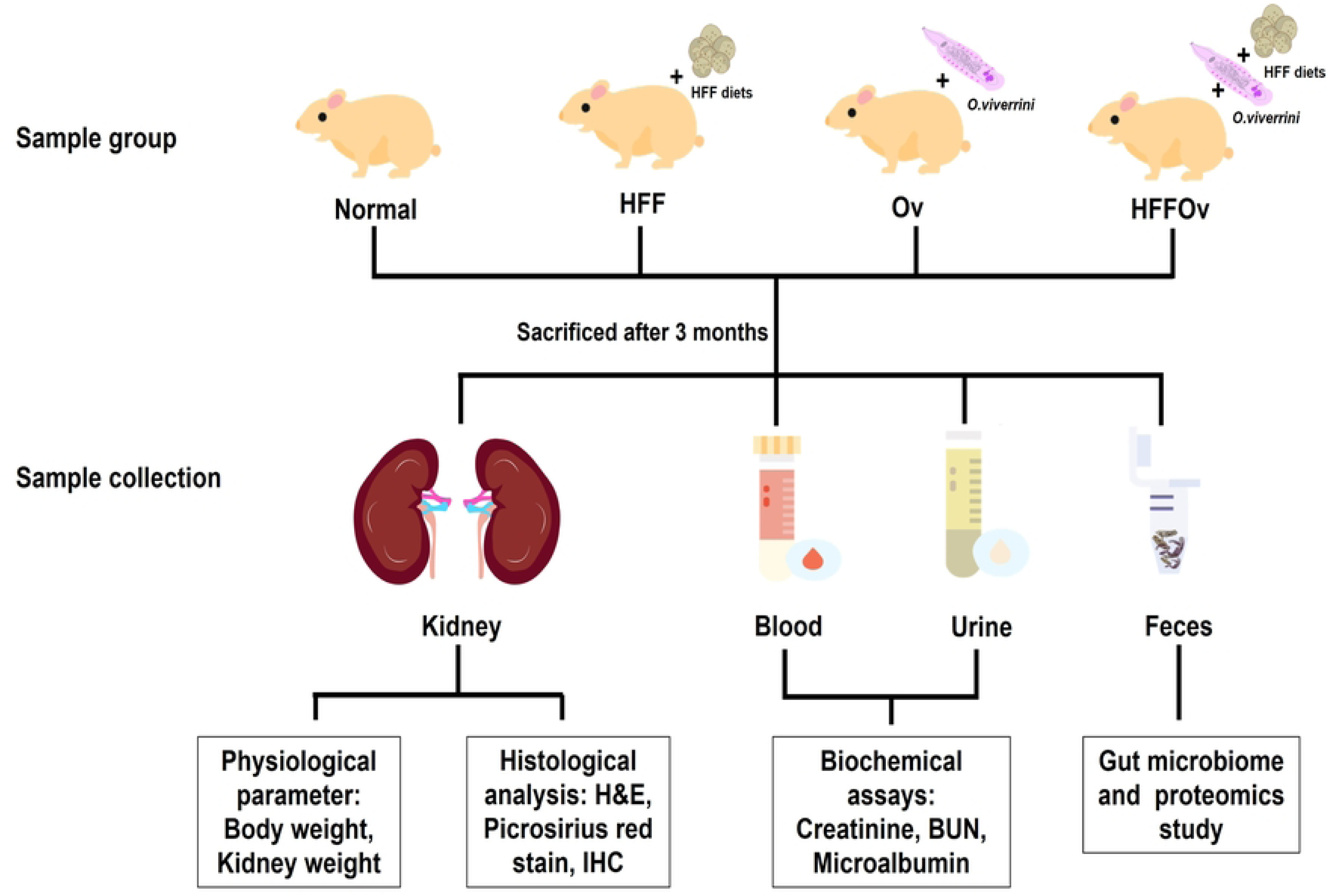
Experimental design. Abbreviations are normal control (Normal: non-fed high fat and high fructose (HFF) diets, and non-infected *Opisthorchis viverrini* (Ov)); hamsters fed only HFF diets; hamsters fed only Ov infection (Ov) and hamsters fed HFF diets and infected with Ov (HFFOv).

### Measurement of serum/urine biochemistry parameters

Blood and urine biochemical parameters of kidney function including blood urea nitrogen (BUN), serum creatinine (SCr) and urine albumin to creatinine ratio were measured after treatments by using the automated analyzers (Cobas 8000 Modular Analysis Series, Roche Diagnostics, Manheim, Germany) at the clinical laboratory of Srinagarind Hospital, Faculty of Medicine, Khon Kaen University, Khon Kaen, Thailand.

### Histopathological of kidney changes investigation

Investigation of the histopathological of kidney changes (tubular damage, inflammatory cell infiltration, and tubular fibrosis), the Mayer’s hematoxylin and eosin (H&E) staining and the picrosirius red methods were used. After fixing with 10% buffered formalin overnight, kidney tissues were cut and embedded with paraffin using automated machine. The paraffin-embedded kidney tissues were cut into 5 um thin sections using microtome. The sections were deparaffinized in xylene and were rehydrated in descending concentrations of ethanol (100%, 95% and 70%, three times for each concentration) and finally in water for 5 minutes each. After rehydration, the slide was stained with Mayer’s hematoxylin for 8 minutes and eosin for 5 minutes and was washed twice with distilled water. The slides were then dehydrated in ascending concentrations of ethanol (70%, 95% and 100%, three times for each concentration) and finally in xylene. Stained slides were mounted with mounting media and air-dried overnight at room temperature. Histopathology was observed under a light microscope.

In the case of histopathological fibrosis investment, the picrosirius red staining method. The tissue section was incubated at 100 °C for 3 minutes and deparaffinized in xylene for 5 minutes (three times) to remove paraffin wax. After following steps, absolute alcohol for 5 minutes (two times), 95% alcohol for 5 minutes (two times), 70% alcohol for 5 minutes, and finally rinsed in the tap water for 5 minutes. After hydration, the slide was stained with Harry’s hematoxylin for 8 minutes and washed in running tap water for 1 minute. The slide was then stained in picrosirius red staining (Solution A for 2 minutes, Solution B for 90 minutes and Solution C for 2 minutes). Slide was then dehydrated via repeat five times in each step as following: 70%, 95% and absolute alcohol, xylene solution and mounted. The appropriate result was stained collagen is red on pale yellow background. The section slides were mounted with mounting solution and air-dried at room temperature for overnight.

### Immunohistochemical study

The immunohistochemical study was performed in kidney sections for protein localization. The paraffin-embedded kidney tissue sections were deparaffinized in xylene and were rehydrated in descending concentrations of ethanol (100%, 95% and 70%, three times for each concentration) and finally in water for 5 minutes each. After rehydration, the antigen was unmasked by autoclaving with sodium citrate buffer (10 mM sodium citrate, 0.05% Tween 20, pH 6.0). Thereafter, slides were immersed in 3% H_2_O_2_ for 10 min for endogenous peroxidase quenching. Then, non-specific binding was blocked by incubating the slides with 5% bovine serum albumin (BSA) for 1 hour at room temperature. The sections were then incubated with primary antibodies diluted in 5% BSA solution at 4 °C for overnight. In the next day, the slides were washed with phosphate buffered saline solution (PBS) followed by incubation with secondary antibody diluted in 5% BSA solution at room temperature for 1 hour. Finally, the immunoreactivity was developed by adding 3,3-diaminobenzidine (DAB) and slides were counterstained with Mayer’s hematoxylin for 2 minutes.

In this study, one primary antibody was utilized: of mouse monoclonal kidney injury molecule-1 (KIM-1) antibody (NBP1-76701, Novus biologicals, CO, USA). The stained sections were examined using a microscope (Carl Zeiss, Oberkochen, Germany). Ten representatives randomly areas at a magnification x200 were monitored. Injury cells scores were analyzed with ImageJ software (National Institutes of Health, Bethesda, MD, USA) [28]

### 16S-rRNA next-generation sequencing analysis

#### Fecal DNA extraction

The fecal of hamster total 19 samples for microbial DNA extraction process was performed using the QIAamp PowerFecal Pro DNA Kit (Qiagen, Hilden, Germany) according to the manufacturer’s instructions. DNA concentration of each sample were measure by Nanodrop2000 spectrophotometer (Thermo Scientific, MA, USA). After extraction and checking, all DNA samples were stored at -20 °C until analysis.

### 16S rRNA gene sequencing and analysis

The V3-V4 region of 16S-rRNA gene was PCR amplified and confirmed using a BioRad C1000TM Thermal Cycler (Biorad, CA, USA). 1.5% gel electrophoresis was used to confirm the size of the product, which should be approximately 450–500 bp.

The 16S rRNA gene was analyzed by sequencing on an Illumina platform (Illumina Inc., CA, USA). The library was prepared by fragmentation of genomic DNA (gDNA) and ligating it to both fragment ends with specific adapters. Paired-end reads were assigned to samples based on their unique barcode [13] and truncation by cutting off the barcode and primer sequences. To generate raw tags paired-end reads which partially overlapped were merged using FLASH (V1.2.7, http://ccb.jhu.edu/software/FLASH/) [29]. Quality filtering on the raw tags was performed under specific filtering conditions to obtain high-quality clean tags [30] according to the Qiime (V1.7.0, http://qiime.org/scripts/split_libraries_fastq.html) quality control process.

Tags were compared to the reference database (Gold database, http://drive5.com/uchime/uchime_download.html) using the UCHIME algorithm (UCHIME Algorithm, http://www.drive5.com/usearch/manual/uchime_algo.html) [31] to detect chimeric sequences, which were then removed [32]. All tags that successfully passed these filtering processes were then analyzed using the Uparse software (Uparse v7.0.1001 http://drive5.com/uparse/) [33]

Sequences sharing ≥97% similarity were assigned to the same Operational Taxonomic Unit (OUT). For each OUT, one representative sequence was analyzed for further annotation. Each representative sequence was checked against the SSU rRNA database of the SILVA database (http://www.arb-silva.de/) [34] using the Mothur software for annotation at each taxonomic rank (threshold:0.8∼1) [35] (kingdom, phylum, class, order, family, and genus).

Information on the abundance of each OTU was normalized relative to the sample with the fewest sequences. Subsequent alpha and beta diversity analyzes were all performed based on these normalized data. Alpha diversity indicates the species diversity of a sample by two indices, including Shannon and Simpson. We calculated these indices using QIIME (version 1.7.0) [36] and plotted the results using R software (version 2.15.3) [37]. Beta diversity analysis was used to evaluate differences among samples in species complexity: the PCoA of Bray-Curtis distance and weighted UniFrac values were calculated using QIIME software (version 1.7.0) [36] for all experimental groups following previous reports [15, 38].

### Meta-Proteomics Analysis of HFFOv group

#### Sample preparation for shotgun proteomics

200 mg of feces were homogenized with 500 µl of 0.5% sodium dodecyl sulfate (SDS) or lysis buffer and then mixed well by vortexed and centrifuged at 10,000g for 15 minutes. All supernatants from each sample were transferred to a new tube, mixed well with 1,000 µl of cold acetone, and incubated overnight at -20 °C. The supernatant was removed from the mixture after centrifugation at 10,000 g for 15 minutes. After precipitation, the protein pellet was dried and stored at -80 °C until analysis.

The Lowry assay determines protein concentration using bovine serum albumin as the standard protein [39]. Used 10 mM dithiothreitol in 10 mM ammonium bicarbonate for breakdown disulfide bonds, and alkylation with 30 mM iodoacetamide in 10 mM ammonium bicarbonate use for prevention of reformation of disulfide bonds in the proteins. The protein samples were digested using sequencing-grade porcine trypsin for 16 h at 37 °C. The tryptic peptides were dried with a speed vacuum concentrator and resuspended in 0.1% formic acid for analysis by nano-liquid chromatography tandem mass spectrometry (nano LC-MS/MS)

#### Liquid chromatography-tandem mass spectrometry (LC/MS–MS)

Tryptic peptide samples were analyzed using the Thermo Scientific Ultimate3000 Nano/Capillary LC System (Thermo Scientific). It was connected to a hybrid quadrupole Q-Tof impact II (Bruker Daltonics; MA, USA) with a nano-captive spray ion source. Peptide digests were packed with Acclaim PepMap RSLC C18, 2 m, 100, and nanoviper after being enriched on a pre-column 300 m i.d. X 5 mm C18 Pepmap 100, 5 m, 100 A (Thermo Scientific). The C18 column was surrounded by a thermostated column oven set at 60 °C. On the analytical column, solvents A containing 0.1% formic acid in water and B containing 0.1% formic acid in 80 % acetonitrile were provided.

Peptides were eluted for 30 minutes at a constant flow rate of 0.30 l/min with a gradient of 5-55% solvent B. CaptiveSpray was used for 1.6 kV electrospray ionization. About 50 l/h of nitrogen was employed as the drying gas. Collision-induced-dissociation (CID) product ion mass spectra were obtained using nitrogen gas as the collision gas. Mass spectra in positive ion mode and MS/MS spectra were obtained at 2 Hz over the range (m/z) 150-2200. Based on the m/z value, the collision energy was changed to 10 eV. The LC-MS analysis was performed three times with each sample.

#### Metaproteomic analysis of samples from the HFFOv group

Maxquant 2.1.0.0 was used to quantify and identify host proteome in fecal samples. The Uniprot database, accessed July 18, 2022, was used for this purpose. Label-free quantification with default MaxQuant settings was performed using a maximum of two missed cleavages, a mass tolerance of 0.6 daltons for the main search, trypsin as the digestive enzyme, carbamidomethylation of cysteine as a fixed modification, and oxidation of methionine and acetylation of the protein N-terminus as variable modifications. Peptides with at least seven amino acids and at least one unique peptide were considered for protein identification and used for further data analysis. All proteins were found in at least 50% of the samples in each group. The median number of modifications per peptide was set at 5. The peptides with the median intensity from three injections were determined as spectral data of the total proteins expressed by the hamster host.

At least 50% of the protein intensities were transformed in Microsoft Excel, resulting in protein expression levels for quantification of protein number (protein ID) and analysis of differentially expressed proteins (DEPs) in the host protein. All samples in each group were selected for analysis of highly expressed proteins (HEPs) using the Jvenn viewer. MetaboAnalyst version 5.0 is a comprehensive, freely available web-based metabolomics analysis platform (https://www.metaboanalyst.ca/, accessed September 14, 2023) used for proteomics analysis. All significant proteins were considered unique and used for the Gene Ontology study by matching the unique protein ID with the Uniprot database. The database STITCH, version 5, was used to predict functional interaction networks between identified proteins and biomolecules and to generate highest confidence (0.900) numbers (http://stitch.embl.de/, accessed September 18, 2023).

### Statistical Analysis

Statistical analyzes were performed using SPSS version 29, GraphPad Prism version 8.4, and Excel. All data were expressed as mean ± SD. Analysis of variance (ANOVA) and Tukey’s test were performed to test for differences between experimental groups using SPSS 29 software (SPSS Inc, IL, USA) and GraphPad Prism version 8.4 (GraphPad sofwere, MA, USA). The intensity of proteins in each group was analyzed using the median value in Excel (Microsoft, WA, USA).

### Data availability

The raw sequencing data have been deposited on the Mendeley Data accession (doi: 10.17632/8hrf8hsx8d.1). The MS/MS raw data and analysis files have been deposited in the ProteomeXchange Consortium (http://proteomecentral.proteomexchange.org) via the jPOST partner repository (https://jpostdb.org) with the data set identifier JPST002317 and PXD045309 (preview URL for reviewers: https://repository.jpostdb.org/preview/2126542412650078fd5e120, Access key: 7273).

## Results

### Effect of combination of HFF diets with Ov infection on biochemical parameters

Effect of combining HFF diets with *O.viverrini* infection on biochemical parameters In this study, hamsters in all groups did not differ significantly in body weight (BW), kidney weight (KW), and KW /BW ratio. However, the body weight of hamsters in the Ov and HFFOv groups showed an upward trend. Clinical biochemical parameters were analyzed to identify HFFOv-induced changes in renal function and/or pathological features. The results showed that the Ov and HFFOv groups had significantly lower urea nitrogen (BUN) (P < 0.05) and serum creatinine (SCr) levels (P < 0.05) compared with the normal and HFF groups, respectively. While ACR increased significantly in both groups, as shown in **Table 1**.

**Table 1.**
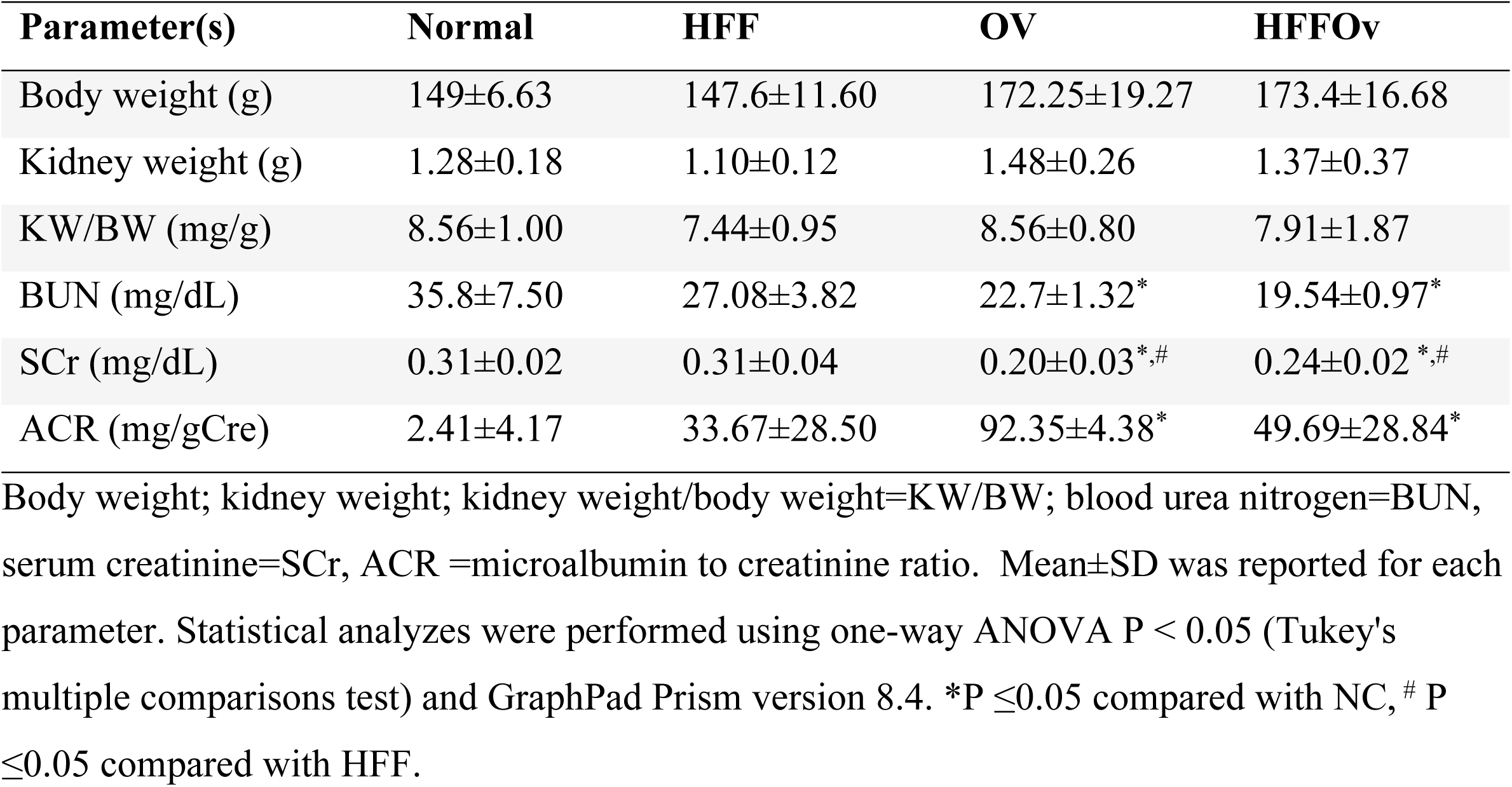
Body weight, Kidney weight, Kidney weight/body weight ratio and biochemical parameters of hamsters.

| Parameter(s) | Normal | HFF | OV | HFFOV |
| --- | --- | --- | --- | --- |
| Body weight (g) | 149±6.63 | 147.6±11.60 | 172.25±19.27 | 173.4±16.68 |
| Kidney weight (g) | 1.28±0.18 | 1.10±0.12 | 1.48±0.26 | 1.37±0.37 |
| KW/BW (mg/g) | 8.56±1.00 | 7.44±0.95 | 8.56±0.80 | 7.91±1.87 |
| BUN (mg/dL) | 35.8±7.50 | 27.08±3.82 | 22.7±1.32* | 19.54±0.97* |
| SCr (mg/dL) | 0.31±0.02 | 0.31±0.04 | 0.20±0.03*,# | 0.24±0.02*,# |
| ACR (mg/gCre) | 2.41±4.17 | 33.67±28.50 | 92.35±4.38* | 49.69±28.84* |

Body weight; kidney weight; kidney weight/body weight=KW/BW; blood urea nitrogen=BUN, serum creatinine=SCr, ACR =microalbumin to creatinine ratio. Mean±SD was reported for each parameter. Statistical analyzes were performed using one-way ANOVA $P < 0.05$ (Tukey's multiple comparisons test) and GraphPad Prism version 8.4. \* $P \leq 0.05$ compared with NC, # $P \leq 0.05$ compared with HFF.

### A combination of HFF diets with *O.viverrini* infection induced kidney fibrosis

Histopathological examination (**Fig 2**) shows representative areas of H&E, KIM -1, and picrosirius red staining in the kidney tissue. H&E staining of the kidney section (**Fig 2A**) clearly showed that there was mild to moderate tubular dilatation along with the appearance of vacuoles in the glomerulus in the HFF and Ov groups compared with the normal group. The HFFOv group showed a higher severity of histopathological changes compared with the HFF and Ov group and the normal group. The HFFOv group had a statistically significant higher positive area of kidney fibrosis as indicated by picrosirius red (**Figs 2B and D**). This result was consistent with the staining of the marker KIM -1 (**Figs 2C and D**), as shown in **Fig 2** (p < 0.05).

**Fig 2.**
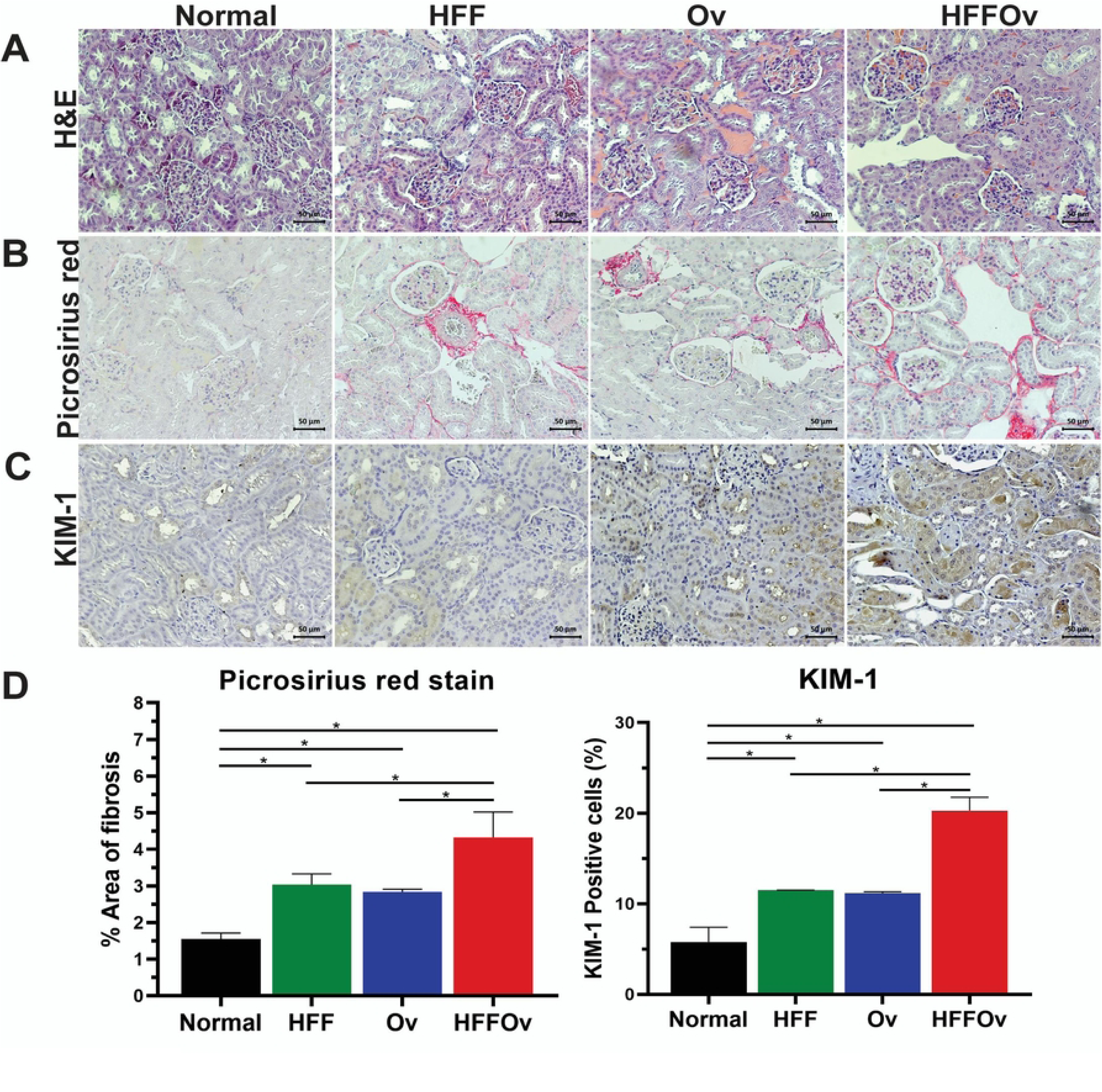
Representative photographs of sections of hamster kidneys. (A) H&E staining (B) picrosirius red staining for fibrosis of kidney tissue and observed under light microscopy 20x, (C) expression of kidney injury molecule-1 (KIM-1) and (D) percentage of picrosirius red staining and KIM-1 positive cells. Results are expressed as mean±SD. Statistical analyses were performed using one-way ANOVA *p< 0.05 (Tukey’s multiple comparisons test) (n = 3 in each group) and GraphPad Prism version 8.4.

### A combination of HFF diets with *O.viverrini* infection disturbs the gut microbiota

A total of high-quality sequences was produced from fecal hamster. The gut microbiome was determined by Illumina Miseq sequencing. The Shannon index, which reflects both the species richness and evenness while the Simpson index reflects only species evenness, was used to determine diversity within experimental groups. The alpha diversity of the HFFOv group was significantly higher than that of the normal group (**Fig 3A**). Principal coordinates analysis (PCoA) based on Bray–Curtis distance and weighted UniFrac distance was used to determine beta diversity between experimental groups (**Figs 3B and C)**. The results showed a clear separation between the groups, indicating that the communities are different in all groups (p= 0.001).

**Fig 3.**
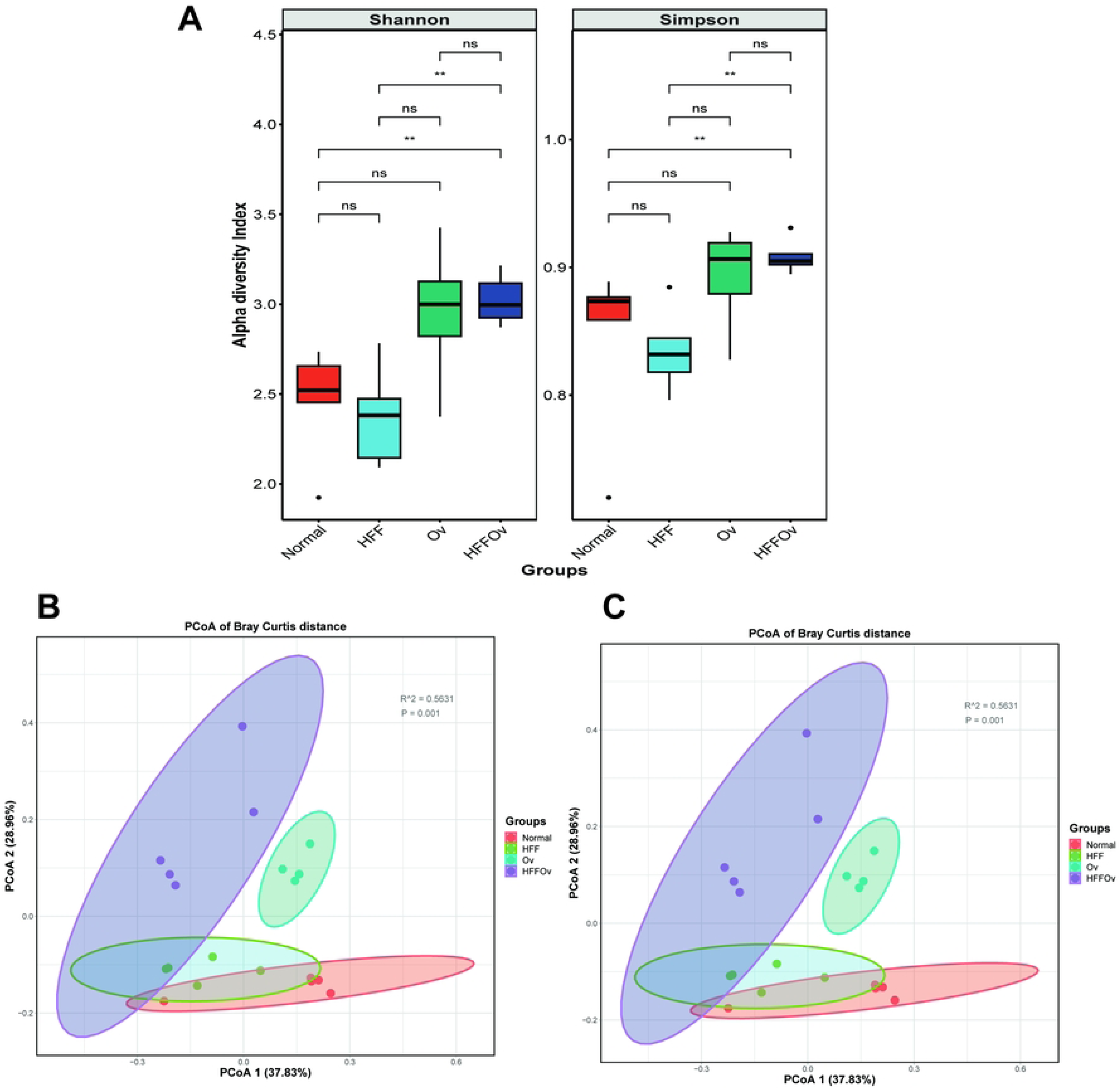
Microbial diversity. (A) Alpha diversity of groups compared on baseline in Normal, HFF, Ov and HFFOv groups presented in boxplots for Shannon and Simpson. Beta diversity of samples in each group of 16S rRNA sequences demonstrates the distribution of samples in orange dots (Normal), green dots (HFF), blue dots (Ov), and purple dots (HFFOv) in (B) PCoA of Bray-Curtis dissimilarity and (C) PCoA of weighted-Unifrac distances with p-values.

At the phylum level (**Figs 4A and B**), *Firmicutes* was the most abundant phylum in all groups. Phylum, *Verrucomicrobia* was most abundant in the HFFOv group, whereas phylum *Actinobacteria* decreased. At the family level (**Figs 4C and D**), *Erypsipelotrichaceae* was the most abundant family in all groups and showed a trend to decrease in the HFFOv group compared to the normal group. Families: *Ruminococaceae, Lachospiraceae, Desulfovibrionaceae* and *Akkermansiaceae* were most abundance in the HFFOv, while *Eggerthellaceae* decreased.

**Fig 4.**
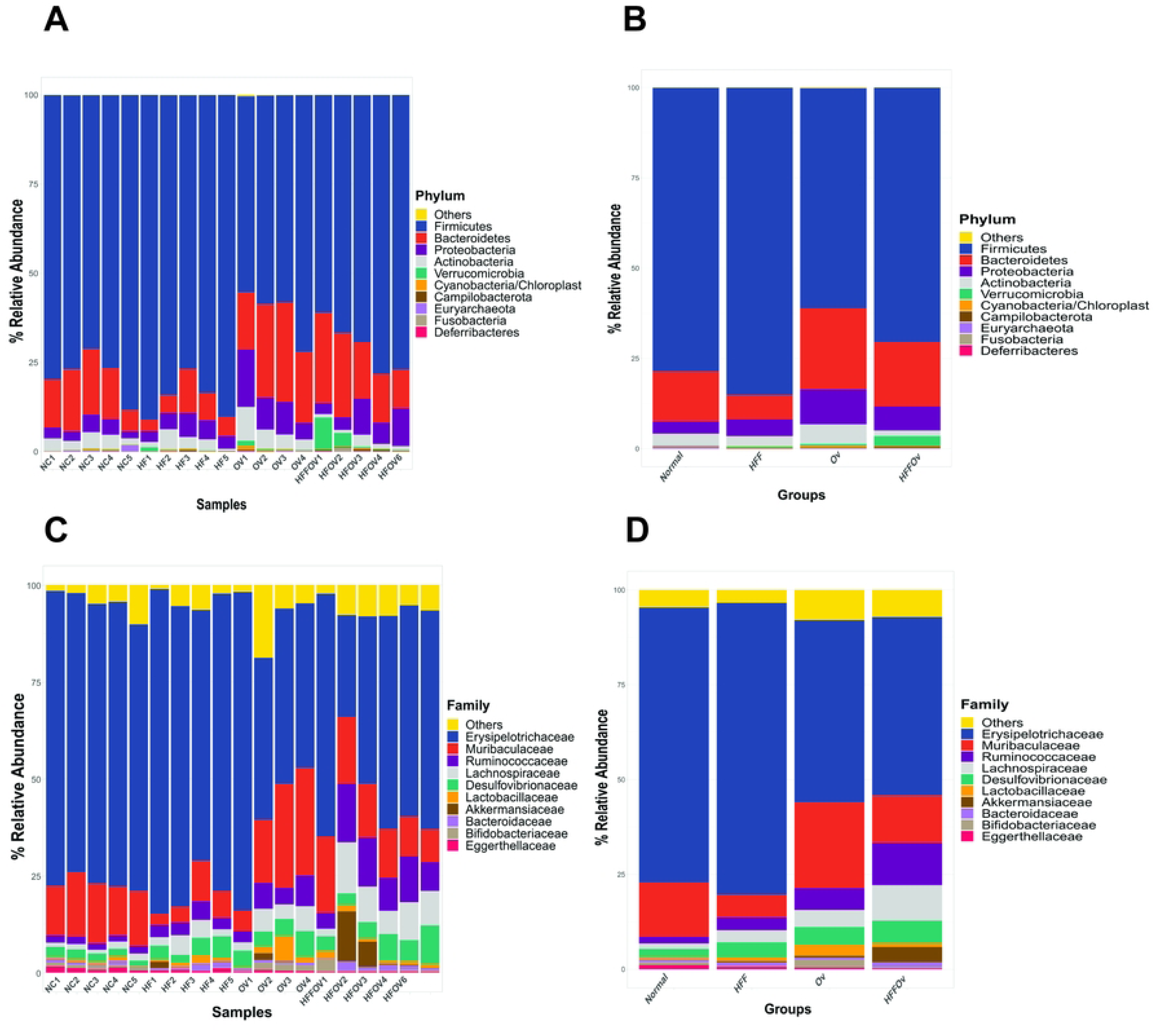
Relative abundance of fecal microbial composition in hamsters of bacterial taxa at the phylum and family levels. Phylum level (A) in each sample and (B) in each group. Family levels (C) in each sample and (D) in each group. Sample name is on the x-axis and relative abundance on the y-axis.

A heatmap of the 35 major families observed in all groups is shown in (**Fig 5).** The results demonstrated that *Eggerthellaceae* and *Methanobacteriaceaein* were most abundant in the normal group compared to HFF, Ov and HFFOv. In contrast, the families *Desulfovibrionaceae, Helicobacteraceae, Selenomonadaceae, Enterobacteriaceae, Streptococcaceae, Prevotellaceae, Clostridiaceae* 1*, Akkermansiaceae, Peptostreptococcaceae, Fusobacteriaceae, Lachnospiraceae, Ruminococcaceae*, and *Porphyromonadaceae* were most abundant in the HFFOv group and decreased significantly in the OV, HFF, and normal groups.

**Fig 5.**
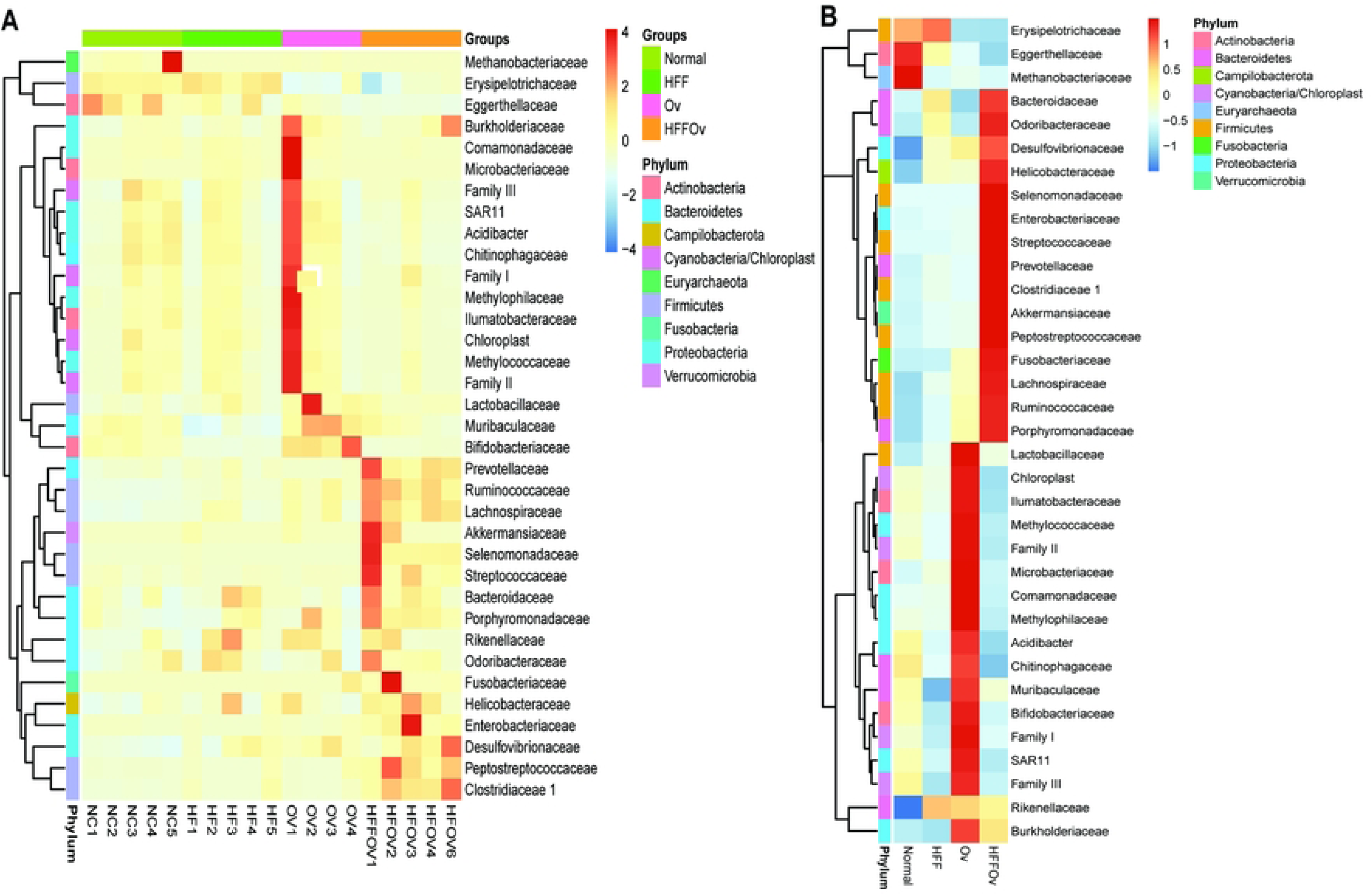
Heat map of the top 35 bacterial families identified in hamster fecal hamster. The abundance of bacterial taxa at the family level (A) in each sample and (B) in each group. Both figures were generated using unsupervised hierarchical cluster analysis (blue, low abundance; red, high abundance).

### Differentially expressed proteins in the HFFOv combination

A total of 14,693 host proteins were identified, of which 243 are significant in all groups, based on MetaboAnalyst software. The 35 most significantly proteins were presented in all groups **(Fig 6A).** In comparison, the group proteins were slightly different **(Fig 6B).** Analysis of differentially expressed proteins (DEPs) using the Jvenn database is shown in (**Fig 6C).** There were 257, 43, 397, and 98 unique proteins identified in the Normal, HFF, Ov, and HFFOv groups respectively. In this study we focus on the HFFOv group to demonstrate that the proteins may be associated with kidney disease. Gene Ontology or functional categories based on Uniprot database were used to analyze the unique proteins in the HFFOv group.

**Fig 6.**
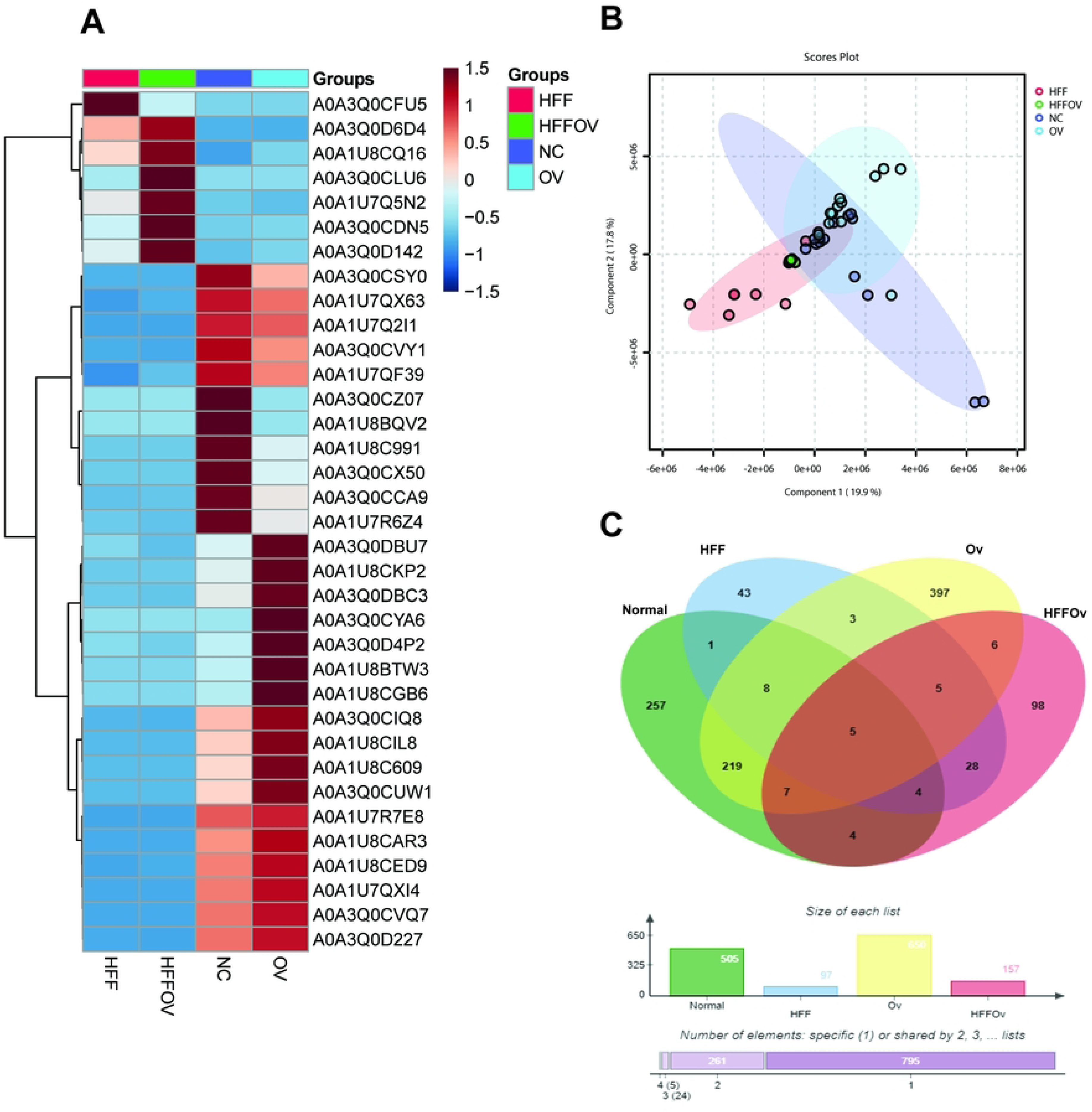
Metaproteomic analysis result. (A) Heatmap of the 35 most abundant host proteins. (B) PLS-DA score plot shows slightly difference between groups. The color of the data: red dots (HFF), green dots (HFFOv), dark blue dots (Ov), and light blue dots (Normal). (C) Venn diagram depicting the number of unique proteins in the HFFOv group.

Gene Ontology or Function categories results show that 46 proteins are concordant and only 15 proteins are associated with kidney structure and/or function, as shown in **Table S1**. Based on the differentially expressed proteins, the highly expressed proteins in the HFFOv group were selected for STITCH analysis. The results showed that TGF-beta-activated kinase 1 and MAP3K7-binding protein 2 (Tab2) interact with biomolecules involved in the inflammatory process, such as trimethylamine N-oxide (TMAO), indoxyl sulfate, p-cresol, indole, and IL -6. In addition, we found that Tab2 interacts with CD14, which is involved in leaky gut (**Fig 7A**).

**Fig 7.**
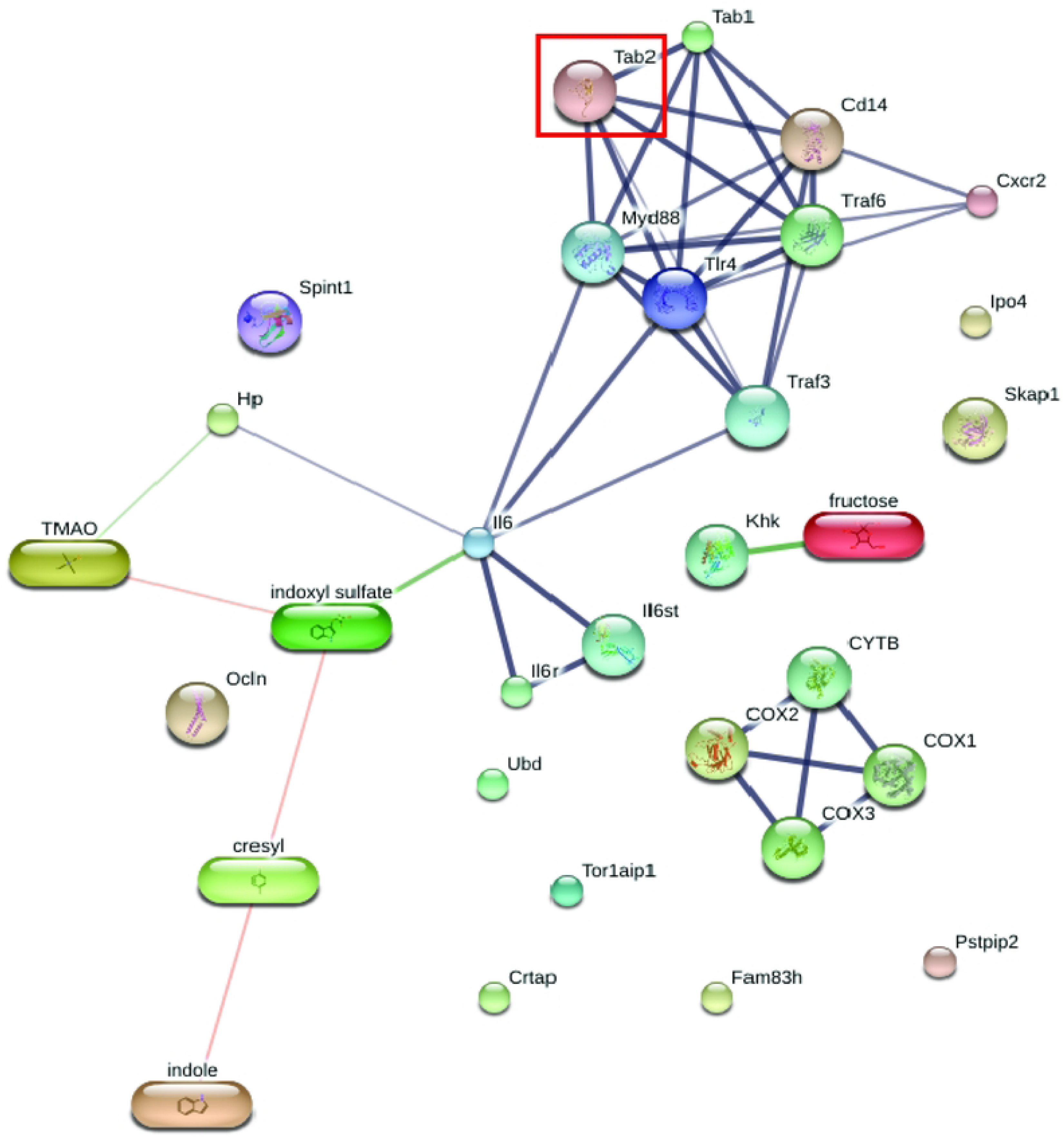
Protein-chemical interactions indicate inflammation and leaky gut. TGF-beta-activated kinase 1 and MAP3K7-binding protein 2 (Tab2) interacted directly with mitogen-activated protein kinase kinase kinase 7 interacting protein 1 (Tab1), TNF receptor-associated factor 6 (Traf6), Toll-like receptor 4 precursor (Tlr4), myeloid differentiation primary response protein MyD88 (Myd88), and monocyte differentiation antigen CD14 (Cd14). The thicker lines represent the highest confidence (0.900). The thin lines represent the medium confidence (0.400).

## Discussion

Ingestion of high fat, high sugar diets was associated with chronic kidney disease (CKD) [3]. We have previously reported that *O. viverrini* infection exacerbates NAFD progression to non-alcoholic steatohepatitis (NASH) in animal models [40], which may be involved in kidney disease [17]. However, the underlying mechanism is unclear. Here, we demonstrated for the first time of alterations in metagenomics and metaproteomics associated with kidney disease in a combination of HFF diets and *O.viverrini* infection in hamster models. We employed the Illumina MiSeq sequencing platform, targeting the V3–V4 region of the 16S rRNA gene and analyzed by LC-MS/MS. The findings revealed that the combination of HFF diets and *O.viverrini* infection promoted the kidney injury by altering the composition of the gut microbiota and host proteome.

First, we investigated the biochemical parameter of renal function, including BUN and SCr. The result shows that both BUN and SCr significantly decreased compared with normal controls. The lower value of BUN may be influenced by the 21 % reduction of dietary protein ingredient. Protein intake affects the levels of BUN and SCr because BUN is a byproduct of protein metabolism [41]. Creatinine is synthesized from creatine, mainly in the liver. Creatine is synthesized from the transamination of the amino acids: arginine, glycine, and methionine [42, 43]. Half of the lost creatinine can be replaced by protein diet [44]. In our study, the nutrient ratio in the diet was adjusted by increasing the fat content. As a result, the protein content of the diet was decreased. For this reason, serum levels BUN and creatinine levels could be lower in the HFF and HFFOv groups due to lower protein intake. The greater decrease in ACR observed in the HFFOv group compared to the other groups may be attributed to the impaired liver function, as this group had more severe liver damage than the other groups [27]. In addition, the lower protein intake in the diet could possibly affect blood albumin levels [45]. Hence, we suggest that these biochemical parameters aren’t appropriate for use as biomarkers of kidney pathology in the HFF diets model.

We examined the histopathology of the kidneys by H&E staining and evaluated tissue fibrosis by picrosirius red staining. The results showed that the combination of HFF diets and *O.viverrini* infection promoted renal injury, such as tubular dilatation, more than *O.viverrini* infection or HFF diets alone. This result is consistent with previous studies showing that showed *O.viverrini* infection and/or HFF are associated with renal injury [10]. This result is supported by significantly higher expression of KIM -1 and fibrosis in the HFFOv group. Elevated levels of KIM -1 have been associated with inflammation and fibrosis in histopathological studies [46]. In addition, renal fibrosis is a reaction process similar to wound healing that occurs in renal injury. If the injury is severe, fibrosis may accumulate in the kidneys, leading to progression of renal dysfunction or eventually renal failure in the future [47].

Several recent studies have demonstrated a strong association between dysbiosis and CKD in both animal models and humans [48–51]. We have demonstrated the effects of changes in the gut in our animal model. The result shows that the combination of HFF diets and *O.viverrini* infection can alter the composition of gut microbiota in hamsters. The alpha diversity in the combination group (HFFOv) was higher than that of the HFF diets alone in our animal model. *O.viverrini* infection might modulate the gut alteration [13, 14]. The results are supported by the improvement of abundance in the HFFOv group, in which bacteria including *Ruminococcus*, *Lachospiraceae*, *Desulfovibrionaceae*, and *Akkermansiaceae* were the most abundant. Similar to many previous reports, the microbiome in the gastrointestinal tract changed: *Lachnospiraceae*, *Ruminococcaceae*, and *Lactobacillaceae* increased, while *Porphyromonadaceae*, *Erysipelotrichaceae*, and *Eubacteriaceae* decreased in hamsters with fluke infestation [52]. Imbalances in the gut microbiota in CKD include changes in both abundance and composition, resulting in increased levels of *Lachnospiraceae*, *Enterobacteriaceae*, and *Ruminococcaceae* [53]. The family *Ruminococcaceae* (genus *Ruminococcus*) is an upward trend in HFFOv group. It can produce pro-inflammatory factor that promote inflammatory bowel syndrome and promote CKD-associated complications, such as glucorhamnan. This molecule induces dendritic cells to produce inflammatory cytokines such as TNF-alpha [54, 55]. In addition, there are also *Lachnospiraceae* and *Akkermansiaceae* that tend to increase HFFOv. Similar to previous studies, *Akkermansiaceae* and *Lachnospiraceae* were regressed in humans on low-protein diet and adenine-induced CKD rats, respectively [56, 57].

Our study determined the host proteome associated with HFF diets associated with *O.viverrini* infection. We focused on the highly expressed proteins (HEPs) in the HFFOv group. We found that TGF-beta-activated kinase 1 (Tab2) proteins were associated with leaky gut (CD14) [58]. A “leaky gut” refers to the impairment of the intestinal barrier, which can be affected by changes in intestinal bacteria and proteins, potentially affecting the permeability of the barrier [53, 59]. Recent studies have linked leaky gut to CKD [60, 61]. In our study, we observed an increase in TGF-beta-activated kinase 1 (Tab2), which is an important cellular signaling molecule that regulates inflammatory processes [62]. Increased inflammation in the gut due to alterations in the microbiome and proteome can lead to disruption of tight junctions in intestinal epithelial cells [63]. Chronic intestinal inflammation may further impair wound healing in the intestine and subsequently increase fibrosis [64]. We also demonstrated the presence of TGF-beta-activated kinase-1 protein, which plays a key role in fibrosis pathogenesis.

Recent research has shown that intestinal fibrosis can affect intestinal structure and function [65]. Consequently, loss of intestinal function due to inflammation and fibrosis may result in bacterial compounds and uremic toxins entering the bloodstream [66]. Gut-derived uremic toxins, including indoxyl sulfate, p-cresyl sulfate, and trimethylamine N-oxide (TMAO), can promote renal inflammation [67]. In addition, TMAO has been associated with increased tubulointerstitial fibrosis and progressive renal dysfunction [68]. Collectively, these findings suggest that the combination of HFF diets with *O.viverrini* infection is associated with renal disease through its effects on intestinal barrier integrity.

We acknowledge to the limitation that our study was conducted in hamsters model. This may affect to the full name of gene, protein data bases for microbiota and host proteome discovery, suggesting further investigation in the patients is required to understand of this change. Furthermore, we performed in small sample size, a larger sample size could increase the power of this finding.

## Conclusion

The combination of HFF diets and chronic opisthorchiasis may promote kidney injury via the alterations of gut microbiome and host proteome in hamsters as shown in **Fig 8**.

**Fig 8.**
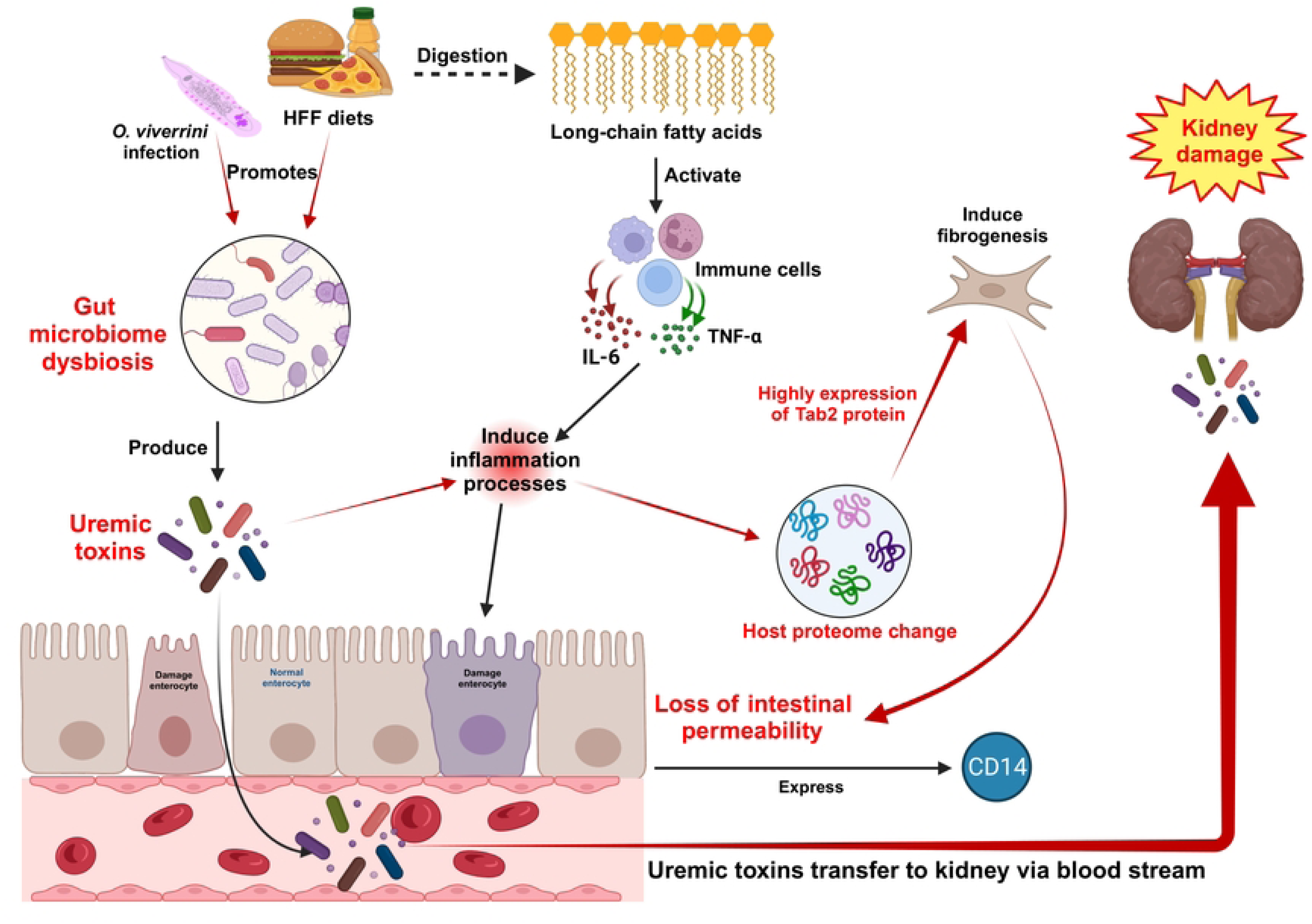
Schematic mechanism of the combination of HFF diets and *O.viverrini* infection promoting the renal injury by alterations the composition of the gut microbiota and host proteome. *Opisthorchis viverrini* infection causes gut dysbiosis and produces uremic toxin, leading to induce inflammation and leaky gut. Ingestion of HFF diets activates immune cells and recruits inflammatory cells, that contribute to intestinal permeability via secretion of IL-6 and TNFα. Loss of intestinal permeability increases the expression of CD14. The host proteome such as TGF-beta-activated kinase 1 and MAP3K7-binding protein 2 (Tab2) is increased during the inflammatory response, which contribute to the activation of fibrogenesis. In addition, uremic toxin leaks through the bloodstream, resulting in kidney damage. The image was created with BioRender.com (license number NM25VIFPY2).

Metagenomics and metaproteomics revealed the highly expressed of TGF-beta-activated kinase 1 (Tab2) proteins and leaky gut (CD14), suggesting that HFF diets and Ov fed might induce gut-derived uremic toxins and leaky gut, eventually to kidney injury/or pathology. We suggest that chronic opisthorchiasis and HFF diets consumption promotes kidney disease via the alterations of metagenomics and metaproteomics. The finding of this study may be an effective strategy for earlier treatment and mitigation of the CKD progression beyond the early stages.

## Supporting information

**S1 Table. Gene ontology proteins associated with kidney disease in feces of hamster in HFFOv group.**

(PDF)

## Acknowledgements

This research was supported by the Fundamental Fund of Khon Kaen University (FF65), which received funding from The National Science Research and Innovation Fund (NSRF). K.T. thanks the scholarship under the Master degree program, Research Fund for Supporting Lecturer to Admit High Potential Student to Study and Research on His Expert Program Year 2022 (651H112), Graduate school, Khon Kaen University, Thailand. We would like to acknowledge Prof. David Blair for editing the manuscript via publication clinic, KKU.

## Author Contributions

**Conceptualization:** Keerapach Tunbenjasiri, Thasanapong Pongking, Chutima Sitthirach, Suppakrit Kongsintaweesuk, Somchai Pinlaor, Porntip Pinlaor

**Data Curation:** Keerapach Tunbenjasiri, Thasanapong Pongking,

**Formal Analysis:** Keerapach Tunbenjasiri, Porntip Pinlaor

**Funding Acquisition:** Porntip Pinlaor

**Investigation:** Keerapach Tunbenjasiri, Chutima Sitthirach, Suppakrit Kongsintaweesuk

**Methodology:** Keerapach Tunbenjasiri, Thasanapong Pongking, Chutima Sitthirach, Suppakrit, Sitiruk Roytrakul, Sawanya Charoenlappanit, Chalongchai Chalermwat, Porntip Pinlaor

**Project Administration:** Keerapach Tunbenjasiri, Somchai Pinlaor, Porntip Pinlaor

**Resources:** Keerapach Tunbenjasiri, Suppakrit Kongsintaweesuk

**Software:** Keerapach Tunbenjasiri, Thasanapong Pongking, Sitiruk Roytrakul, Sawanya Charoenlappanit, Sirinapha Klungsaeng

**Supervision:** Porntip Pinlaor

**Validation:** Keerapach Tunbenjasiri, Thasanapong Pongking, Suppakrit Kongsintaweesuk, Somchai Pinlaor, Porntip Pinlaor

**Visualization:** Keerapach Tunbenjasiri, Porntip Pinlaor

**Writing – Original Draft Preparation:** Keerapach Tunbenjasiri, Porntip Pinlaor

**Writing – Review & Editing:** Keerapach Tunbenjasiri, Thasanapong Pongking, Chutima Sitthirach, Suppakrit Kongsintaweesuk, Sitiruk Roytrakul, Sawanya Charoenlappanit, Sirinapha Klungsaeng, Sirirat Anutrakulchai, Chalongchai Chalermwat, Somchai Pinlaor, Porntip Pinlaor

